# Effect of Histidine Covalent Modification on Strigolactone Receptor Activation and Selectivity

**DOI:** 10.1101/2022.07.13.499796

**Authors:** Jiming Chen, Diwakar Shukla

**Affiliations:** Department of Chemical and Biomolecular Engineering, University of Illinois Urbana-Champaign, Urbana, IL, 61801, United States; Center for Biophysics and Quantitative Biology, University of Illinois Urbana-Champaign, Urbana, IL, 61801, United States; Department of Plant Biology, University of Illinois Urbana-Champaign, Urbana, IL 61801, United States; Department of Bioengineering, University of Illinois Urbana-Champaign, Urbana, IL 61801, United States

## Abstract

The parasitic weed *Striga* has led to billions of dollars’ worth of agricultural productivity loss worldwide. *Striga* detects host plants using the plant hormone strigolactone. Early steps in the strigolactone signaling pathway involve substrate binding and hydrolysis followed by a conformational change to an “active” or “closed” state, after which it associates with a MAX2-family downstream signaling partner. The structures of the inactive and active states of strigolactone receptors are known through X-ray crystallography, and the transition pathway of from the inactive to active state in *apo* receptors has previously been characterized using molecular dynamics simulations. However, it also has been suggested that a covalent butenolide modification of the receptor on the catalytic histidine through substrate hydrolysis promotes formation of the active state. Using molecular dynamics simulations, we show that the presence of the covalent butenolide enhances activation in both *At* D14 and *Sh*HTL7, but the enhancement is ∼50 times greater in *Sh*HTL7. We also show that several conserved interactions with the covalent butenolide modification promote transition to the active state in both *At* D14 (non-parasite) and *Sh*HTL7 (parasite). Finally, we demonstrate that the enhanced activation of *Sh*HTL7 likely results from disruption of *Sh*HTL7-specific histidine interactions that inhibited activation in the *apo* case.

## Introduction

Strigolactone signaling in plants is responsible for regulating shoot branching, root architecture, and hypocotyl elongation. ^1–3^ They have also been found to stimulate germination in parasitic plants.^4^ The canonical model for the early steps of strigolactone signaling are binding of the hormone to the receptor, substrate hydrolysis leading to covalent modification to the receptor by a fragment of the ligand, conformational change to the “closed” or “active” state, and association of the receptor in its active state to MAX2 and SMXL signalling partners.^5,6^ This Skp-Cullin-F-box (SCF) complex is then degraded, leading to downstream signaling effects. ^5–9^ Despite 78% sequence similarity and highly conserved structure, one strigolactone receptor found in witchweed *Sh*HTL7, uniquely exhibits picomolar sensitivity to strigolactone in inducing a downstream signaling response. ^10^ Wang *et al*. attributed this difference primarily to uniquely high MAX2 affinity exhibited by *Sh*HTL7, leading to enhanced stability of the signaling complex. ^11^ However, since the signaling complex between D14/HTL proteins and MAX2 contains D14/HTL in its active state, it is also possible that higher propensity to undergo conformational transition to the active state in response to strigolactone binding and hydrolysis may lead to enhanced MAX2 affinity and thus greater signaling activity.

In our previous work, we characterized the activation mechanisms of *apo At* D14 and *Sh*HTL7 in the absence of substrate and determined that while both receptors are able to undergo constitutive activation, several key interactions with the catalytic histidine selectively inhibit *apo* receptor activation in *Sh*HTL7, which has lower constitutive activity than *At* D14.^15^ Furthermore, a D218A mutant of *At* D14 in which the interaction between H247 and D218 is disrupted has been shown experimentally to induce a signalling response in the absence of substrate. ^16^ Since this mutant is catalytically inactive, this result suggests that strigolactone hydrolysis is not necessary for receptor activation, despite other results indicating that covalent modification of the histidine induced by strigolactone hydrolysis promotes strigolactone signaling. ^5,7^ Our computational characterization of this mutant indicates that it undergoes *apo* activation at a higher rate than the wild-type. ^15^ These results indicate that if the catalytic histidine is covalently modified, disruptions in these interactions could enhance strigolactone receptor activation. Furthermore, since these H246 interactions selectively inhibit activation in *Sh*HTL7 over *At* D14, introduction of a covalent modification could have a greater influence on *Sh*HTL7 activation than *At* D14 activation. In light of these findings, two outstanding questions are (i) how covalent modification to the catalytic histidine affects the propensity for activation, and (ii) how this response differs between *At* D14 and *Sh*HTL7. Here, we evaluate the effect of a histidine-covalent butenolide ring (D-ring) on the activation process of *At* D14 and *Sh*HTL7. The active and inactive states of *At* D14 are shown in Fig. 1a-b, and the structure of the covalent D-ring bound to histidine is show in Fig. 1c-d.

**Fig. 1.**
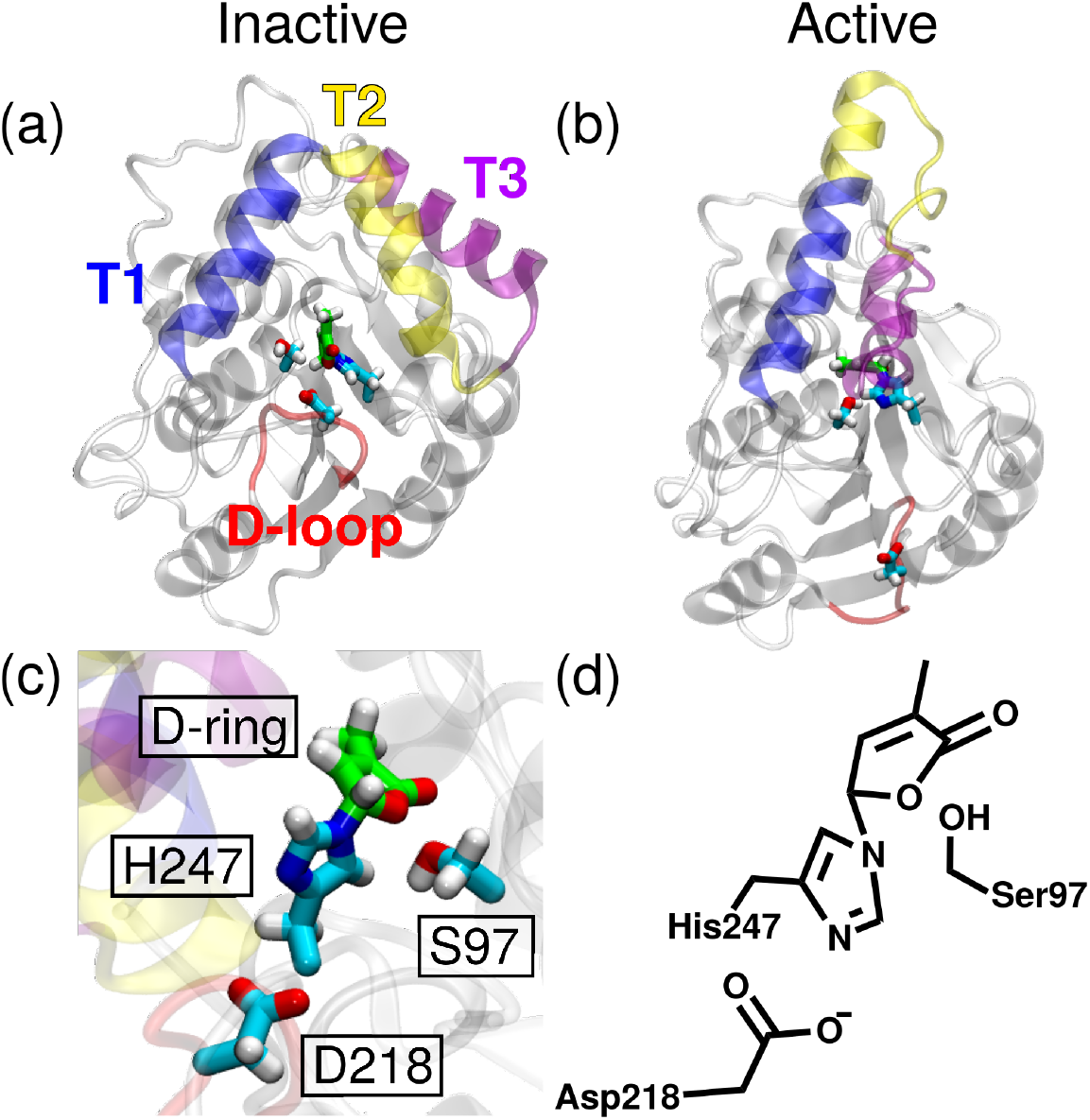
(a) Inactive and (b) active states of *At* D14. The T1 (residues E138N151), T2 (residues N151G165), and T3 (residues P169M182) helices are shown in blue, yellow, and purple, respectively, and the aspartate loop (D-loop, residues A216A223) is shown in red. The inactive state was sourced from PDB code 4IH4, ^12^ and the active state was sourced from PDB code 5HZG^5^ with missing D-loop residues added using Modeller.^13,14^ (c) The covalent modification to the catalytic histidine, and (d) a line drawing of the catalytic triad with the covalent modification on histidine.

To characterize the activation mechanisms of covalently-modified *At* D14 and *Sh*HTL7, we employed long-timescale (*>*1 ms aggregate) molecular dynamics (MD) simulations. Molecular dynamics simulations have previously been used to study effects of other covalent modifications on proteins,^17–19^ including other plant hormone receptors.^20–22^ To extract thermodynamic and kinetic information from these simulation datasets, we constructed Markov state models.^23,24^ Briefly, these models extract kinetic and thermodynamic information from a set of simulation data by discretizing the data into states and estimating transition probabilities between states. This method has previously been applied to study protein conformational dynamics and substrate interactions for several other plant hormone systems. ^21,25–27^ Using these methods, we demonstrate that presence of a substrate hydrolysis-induced covalent modification promotes receptor activation in both *At* D14 and *Sh*HTL7. Furthermore, the enhancement of activation is greater for *Sh*HTL7 than *At* D14, suggesting *Sh*HTL7 is more responsive to the presence of substrate than *At* D14. This indicates that greater enhancement of activation in response to covalent modification plays a significant role in producing the high signaling activity observed in *Sh*HTL7.

## Results

### Free energy landscapes show enhancement of activation by covalent D-ring

In our previous work, we identified four molecular switches that determine whether a receptor is in its active or inactive state by comparing the inactive and active state crystal structures: closure of the T1 and T3 helices, partial unfolding of the T2 helix, elongation of the T1 helix with a portion of the T2 helix, and detachment of the D-loop.^15^ Using the same molecular switches, we computed free energy landscapes of activation for the covalently modified receptors (Fig. 2) using the relation *F* = −*RT* ln *P*, where *P* is a probability density. From these landscapes, the relative stabilities of the active and inactive states are similar (within ∼1 kcal/mol of each other) in both *At* D14 and *Sh*HTL7 with the covalent D-ring present. In the *apo* case, the active state of *At* D14 was ∼ 1-2 kcal/mol higher in free energy than the inactive state, and the active state of *Sh*HTL7 was ∼3 kcal/mol higher in free energy than the inactive state. ^15^ This indicates that while formation of the active state is promoted by the covalent D-ring in both *At* D14 and *Sh*HTL7, the modification enhances activation to a greater extent in *Sh*HTL7 than in *At* D14.

**Fig. 2.**
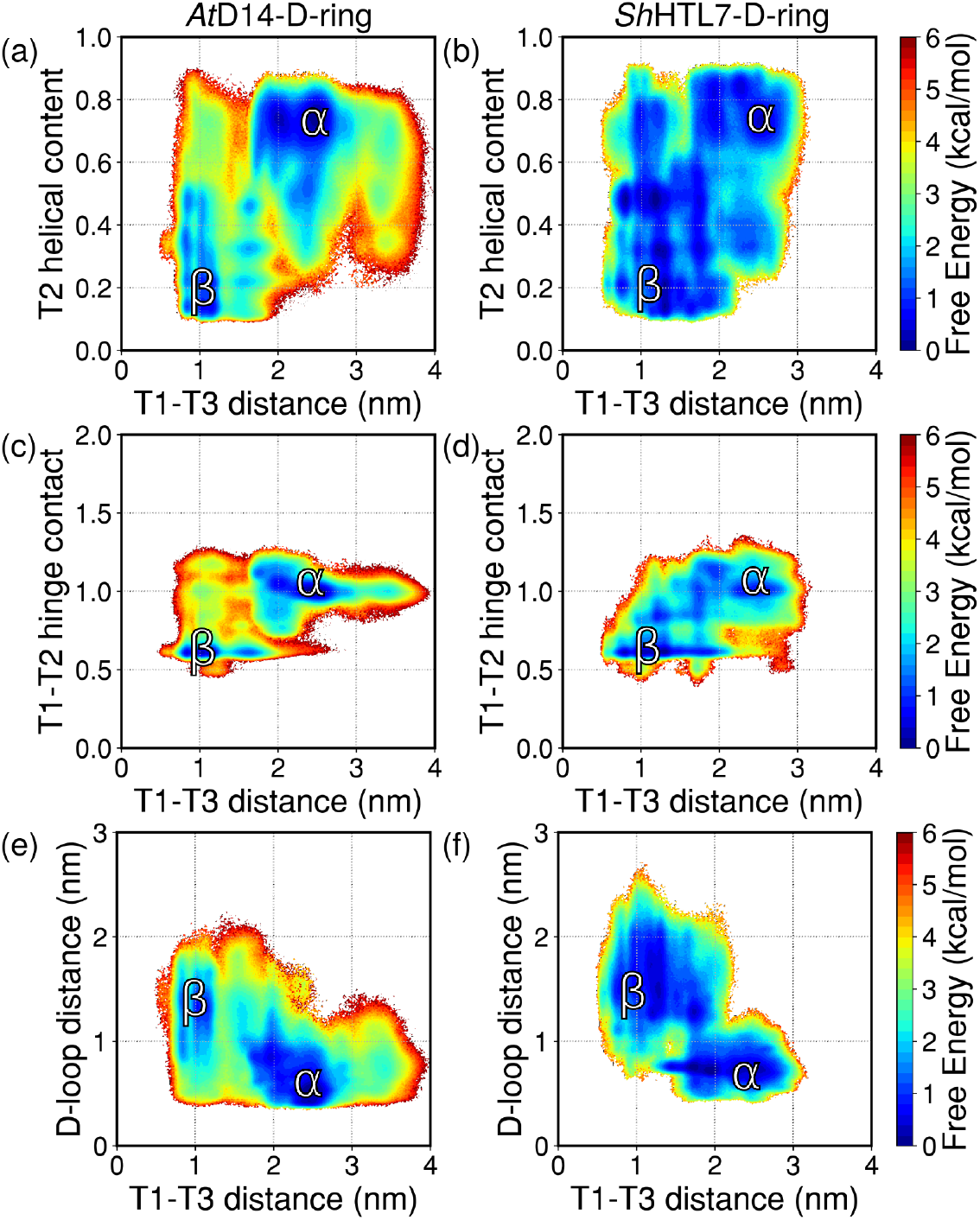
Free energy landscapes of the activation process in covalent D-ring modified *At* D14 (a,c,e) and *Sh*HTL7 (b,d,f). These landscapes show the activation process projected onto the T1-T3 helix distance and the other molecular swtiches: unfolding of the T2 helix (a, b), extension of the T1 helix (c, d), and D-loop detachment (e, f). The inactive state in each landscape is labeled as *α* and the active state is labeled as *β*.

To estimate the free energy barrier of transition between the inactive and active states, we performed time-lagged independent component analysis (TICA) on each dataset.^23^ Briefly, TICA identifies a linear combination of a given set of structural metrics that correspond to the slowest process found in the simulations, which are associated with the highest free energy barriers. Free energy landscapes of the activation pathway projected onto TICA coordinates are shown in Fig. S1. Based on the TICA landscapes, the free energy barrier of the activation process is ∼4 kcal/mol in *At* D14 and ∼2-3 kcal/mol in *Sh*HTL7. These are both decreases from the ∼5 kcal/mol barrier seen in both *apo At* D14 and *Sh*HTL7. ^15^ Again, these values show that while presence of the covalent D-ring enhances activation in both *At* D14 and *Sh*HTL7, the enhancement is greater in *Sh*HTL7. These differences in free energies indicate that the activation enhancements in the two proteins differ by approximately an order of magnitude. To quantify the enhancement of active state formation further, we also compute probabilities of activation for covalent D-ring-modified *At* D14 and *Sh*HTL7.

### Activation probabilities show 50-fold greater activation enhancement in *Sh* HTL7 than *At* D14

To evaluate the enhancement of activation by the covalent D-ring further, we computed equilibrium probabilities of each of the four molecular switches (Fig 3). The active definitions of each molecular switch follow the same definitions as used in our previous work: T1T3 distance *<* 1.3 nm, T2 helical content *<* 0.6, T1T2 hinge contact *<* 0.75 nm, and D-loop distance *>* 1.0 nm.^15^ Overall activation probability was computed as the product of each individual molecular switch. These overall probabilities indicate that presence of the covalent D-ring leads to a ∼2-fold increase in presence of the active state in *At* D14 and a ∼100-fold increase in presence of the active state in *Sh*HTL7 (Fig. 4). Calculated values of molecular switch probabilities and overall activation probabilities are shown in Table S1. In *At* D14, this enhancement is largely driven by the increase in T2 helix unfolding, with a ∼1.8-fold enhancement of this molecular switch activation probability. In contrast, the presence of the covalent D-ring in *Sh*HTL7 increases each of the molecular switches at least 2-fold, with the greatest enhancement seen in the nearly 10-fold increase in probability of the T1-T3 helices being close together. These probabilities are able to quantify the enhancement of molecular switch activation in both proteins, however, further investigation into how the molecular switches modulate the conformational change between the inactive and active states requires investigation of the transition pathways.

**Fig. 3.**
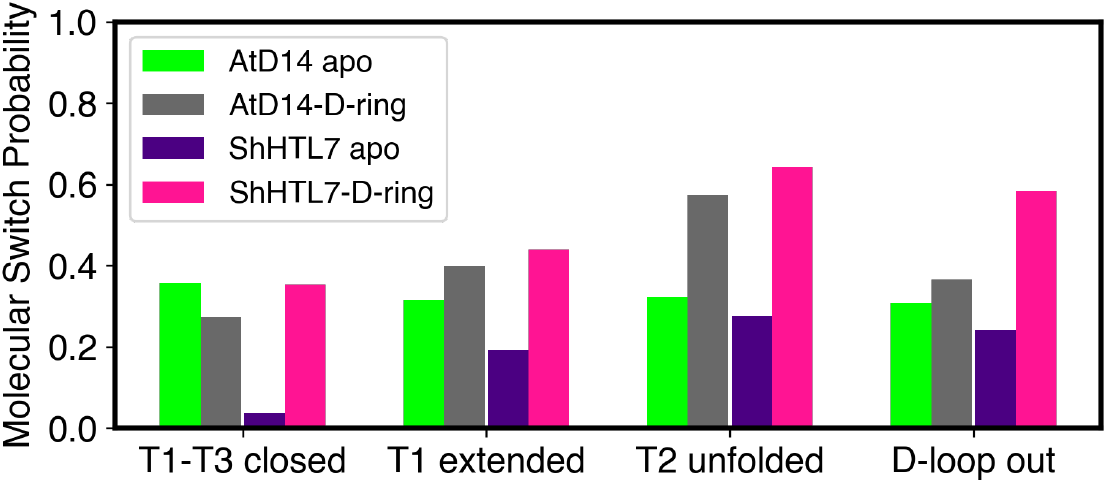
Molecular switch probabilities for *apo* and covalent D-ring modified *At* D14 and *Sh*HTL7. These probabilities show modest increases in covalent D-ring modified *At* D14 (with a slight decrease in T1-T3 closed probability) and larger increases in covalent D-ring modified *Sh*HTL7.

**Fig. 4.**
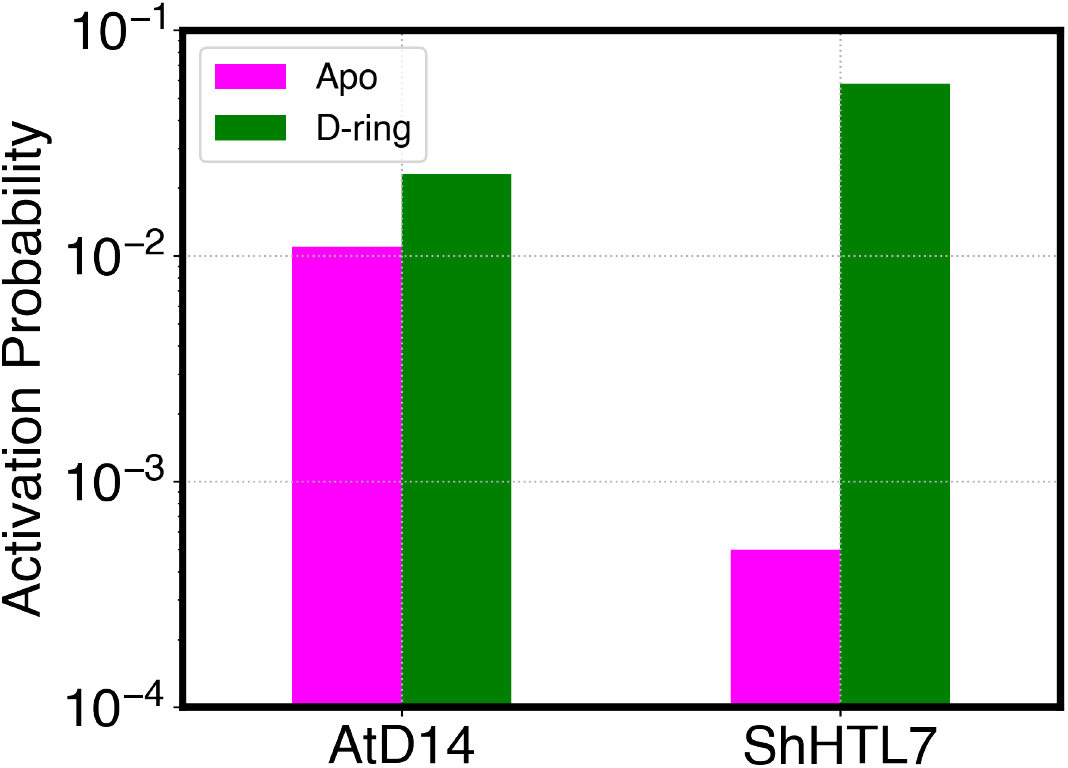
Overall activation probabilities for *apo* and covalent D-ring modified *At* D14 and *Sh*HTL7. These probabilities show a modest (∼2-fold overall) enhancement of activation by the covalent modification in *At* D14 and a large (∼100-fold) enhancement in *Sh*HTL7.

### Transition paths of activation show T1-T3 pocket closure as main barrier to activation

To determine the transition pathways between the inactive and active states of *At* D14 and *Sh*HTL7, we employed transition path theory. ^28^ Top pathways accounting for 99% of the flux between inactive and active states are shown in Fig. 5 and are labeled with Roman numerals in decreasing order of flux. As was observed in the *apo* case,^15^ both D-ring-modified *At* D14 and *Sh*HTL7 have several intermediates along their activation pathways in which multiple molecular switches are in partially active states. This indicates that there is a high degree of coupling between molecular switches. Similarly to the *apo* case, the last molecular switch to activate fully in both protein is the closure of the pocket via merging of the T1 and T3 helices, indicating that T1-T3 closure remains a key barrier to activation. However, in the *apo* case, the detachment of the D-loop was also a key barrier to activation, and its full activation occured in tandem with T1-T3 closure. For the covalent D-ring modified systems, free energy landscapes projected onto the D-loop molecular switch (Fig. 2e-f), and the top pathways from TPT show that the D-loop molecular switch becomes fully active prior to full T1-T3 closure (Fig. 5). These indicate that presence of the covalent D-ring lowers the free energy barrier for D-loop detachment. In light of the consistent observation that the covalent D-ring enhances receptor activation, we investigated the contacts formed by the covalent D-ring to determine the mechanism by which it enhances activation.

**Fig. 5.**
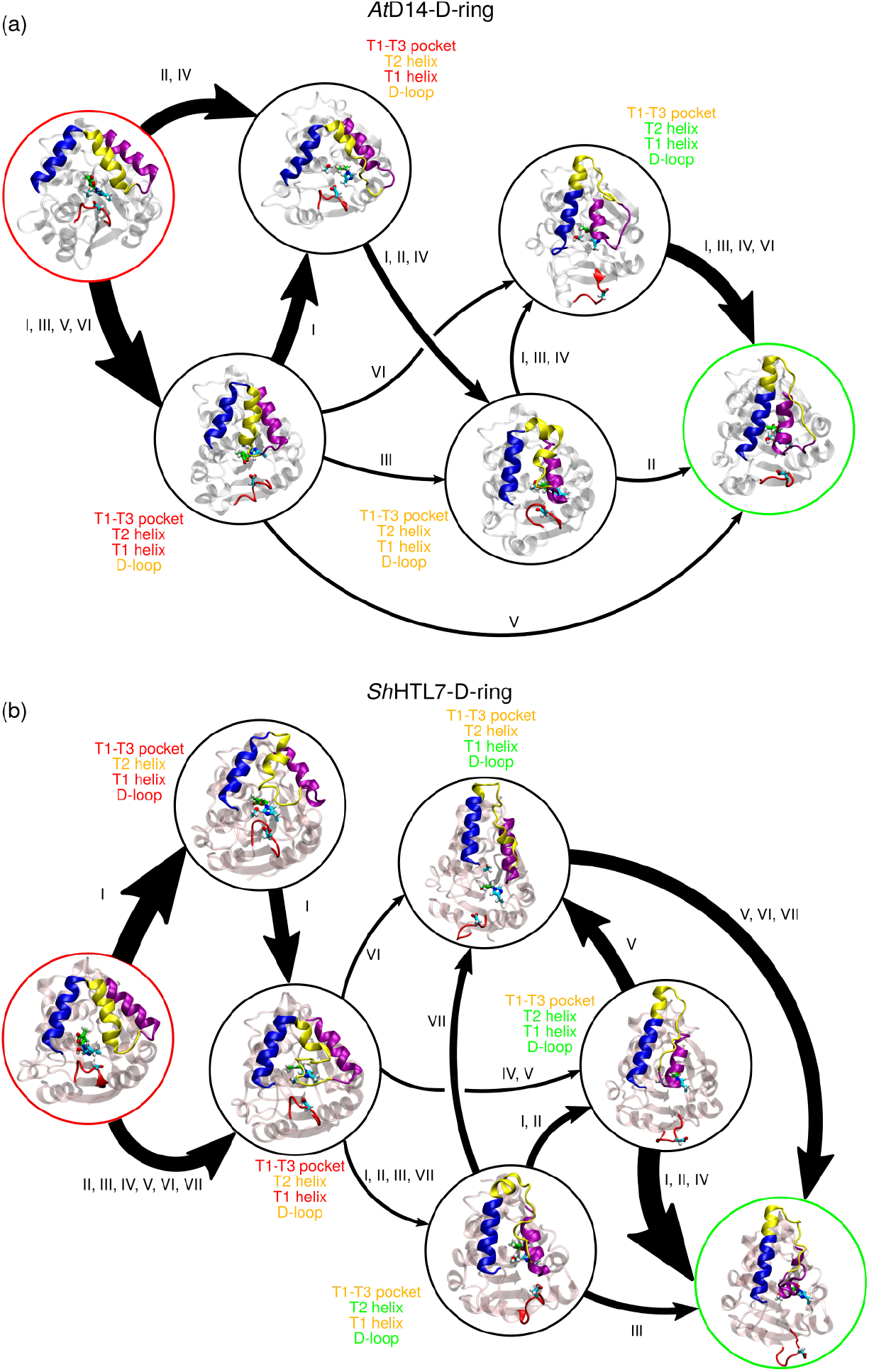
Major transition paths for covalent D-ring modified (a) *At* D14 and (b) *Sh*HTL7. Arrow thickness correspond to relative orders of magnitudes of inverse mean first passage time between states. Each intermediate is labeled with the configuration of the four molecular switches, with red, orange, and green indicating inactive, partially active, and active respectively.

### Conserved interactions with covalent D-ring promote formation of active state

To determine the mechanism by which the covalent D-ring promotes receptor activation, we computed the equilibrium contact probabilities of each residue in *At* D14 and *Sh*HTL7 with the covalent D-ring from our simulation data to identify key interactions formed by the covalent D-ring. Contact probabilities of residues with the covalent D-ring and each protein residue are shown in Fig. S2, and the twenty residues in each protein with the highest D-ring contact probability along with their respective contact probabilities are listed in Table S2. The contact probability of the D-ring with H247/246 was found to be 1.0 in both *At* D14 and *Sh*HTL7, which is expected since the D-ring is covalently bound to H247/246. Additionally, the other two catalytic triad residues S97/95 and D218/217 were found to have high contact probability with the D-ring in both proteins, which is expected due to close proximity of these residues with H247/246. Other regions of high contact probability include several residues on and adjacent to the T2-T3 lid helices (shown in yellow and purple in Fig. 1) as well as F28/Y24-T30/T26. For further investigation into the significance of these high-Dring contact regions, we investigated whether D-ring contacts with these residues affect the activation process.

To determine the impact of these D-ring contacts on activation, we computed free energy landscapes projected onto D-ring-residue distances of top D-ring contacting residues and T1-T3 distance. This molecular switch was chosen as a proxy for activation based on TPT results showing that is the final molecular switch to activate (Fig. 5). Based on these landscapes, key interactions that promote activation are between the D-ring and F28/Y24T30/T26, F175/Y174, and T178/T77. Free energy landscapes projected onto T1-T3 and F28/Y24-D-ring distances, T1-T3 and F175/Y174 distances, and T178/T77 distances are shown in Fig. 6a-c, respectively. Landscapes for the other top D-ring contacts that do not exhibit a stabilizing effect on the active state are shown in Fig. S3. These landscapes show that in both *At* D14 and *Sh*HTL7, interaction with the F28/Y24 residue stabilizes the active state (low T1-T3 distance) and lowers the free energy barrier to formation of the active state by ∼1-2kcal/mol (Fig. 6a). This interaction can be formed in both active(lowT1-T3 distance) and inactive-like (high T1-T3 distance) states in both *At* D14 and *Sh*HTL7, indicating that the lowering of the free energy barrier of activation is a key manner in which this contact promotes formation of the active state. The two subsequent residues, G29-T30 in *At* D14 and G27-T28 *Sh*HTL7 also have a similar stabilizing effect on the active state. Free energy landscapes of these interactions are shown in Fig. S4.

**Fig. 6.**
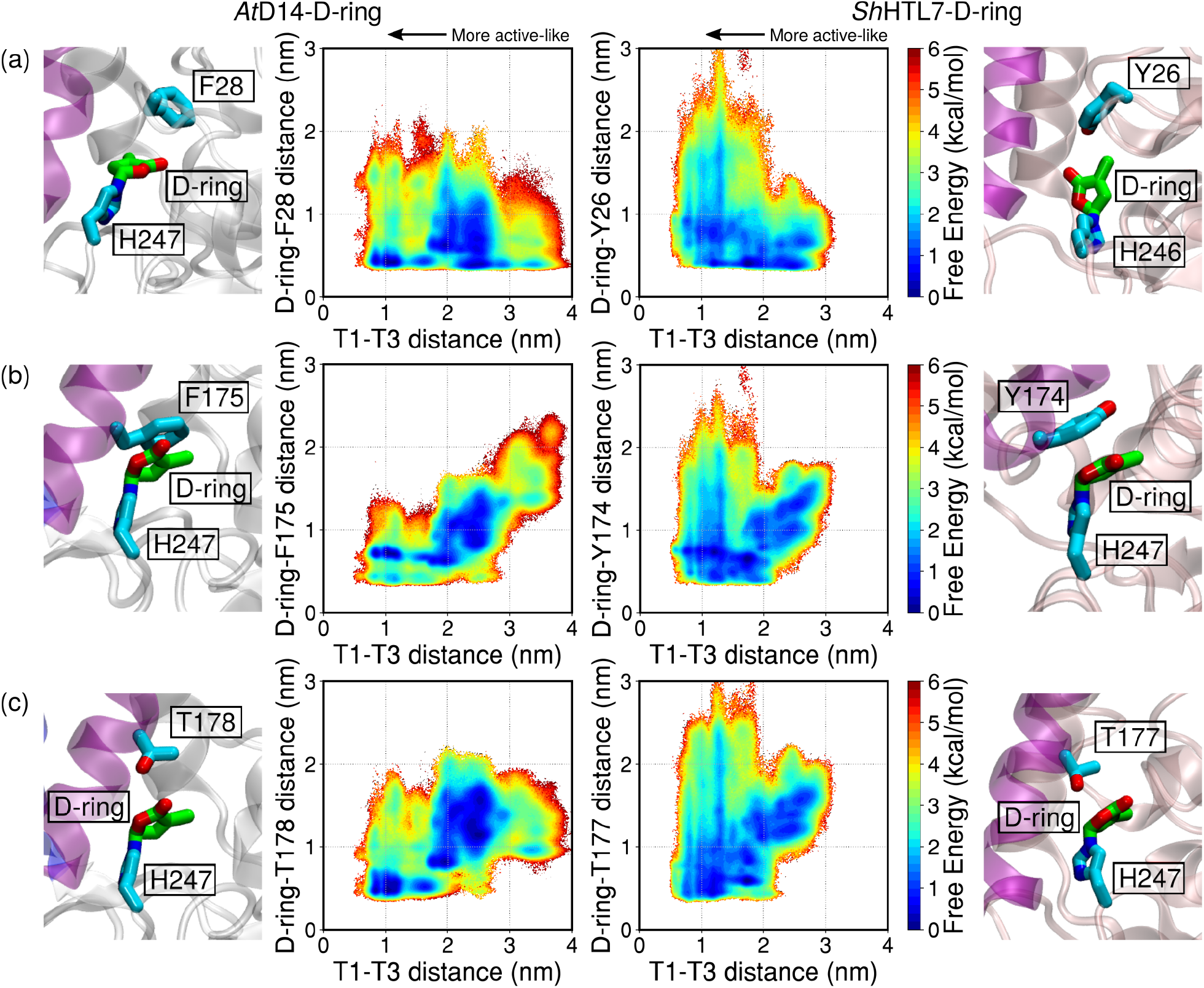
Free energy landscapes and accompanying structural images showing coupling between interactions of the covalent D-ring with high-contact-probability residues and the activation of the T1-T3 closure molecular switch. Key residues identified based on high contact probability are (a) F28 (*At* D14)/Y26 (*Sh*HTL7) and (b) T178 (*At* D14)/T177 (*Sh*HTL7). More active-like states are found in regions with low T1-T3 distance, indicating closure of the binding pocket.

In addition to the F28-T30/G27-T28 residues, two residues on the T3 helix were seen to stabilize the transition to the active state. The covalent D-ring is seen to interact with F175 in *At* D14 and Y174 in *Sh*HTL7. In both proteins, this interaction is seen to lower the barrier of transitioning to the active state as well as stabilize the active state, as shown in (Fig. 6b). Similarly, the covalent D-ring interaction with T178/T77 stabilizes the active state in both *At* D14 and *Sh*HTL7 (Fig. 6c). This interaction does not form in inactive-like states (T1-T3 distance *>* 2.5 nm) in either protein, indicating that this interaction acts as a stabilizing interaction for active-like states. To evaluate the significance of these results further, we performed a sequence conservation analysis of each of the sites F28/Y24T30/T26, F175/Y174, and T178/T77 (Table S3). All of these sites show high conservation among close homologues of *At* D14, *Sh*HTL7, and *At* KAI2 (Table S4). While these conserved stabilizing contacts provide an explanation for why the covalent D-ring enhances activation in both proteins, they do not explain why the enhancement is greater in *Sh*HTL7. To determine the origin of the selectivity in activation enhancement, we investigated the effect of the covalent D-ring on several *Sh*HTL7-specific contacts we identified in our previous work that inhibit *apo* activation in *Sh*HTL7.

### Covalent D-ring disrupts key activation-inhibiting H246 contacts in *Sh* HTL7

In the *apo* state, we identified several *Sh*HTL7-specific interactions with H246 that inhibit the detachment of the D-loop and closure of the pocket by the T1-T3 helices, thus inhibiting activation.^15^ Since H246 is the site of the covalent D-ring modification, we evaluated the influence of the covalent D-ring on these activation-inhibiting interactions in *Sh*HTL7. One interaction that inhibited D-loop detachment was the H246-S168 contact. The presence of this interaction in *apo Sh*HTL7 stabilizes the protein in D-loop-attached conformations such that there was a high free energy barrier for activation of the D-loop molecular switch (Fig. 7a,c). However, in the presence of the covalent D-ring modification, this interaction is disrupted, as shown by stabilization of states in which H246 and S168 are far apart (Fig. 7b,d). Importantly, when this interaction is not present, the free energy barrier to activation of the D-loop molecular switch is lowered from ∼4 kcal/mol to ∼1-2 kcal/mol. This is consistent with previous observation that D-loop detachment is significantly enhanced in the presence of the covalent D-ring (Table 1).

**Fig. 7.**
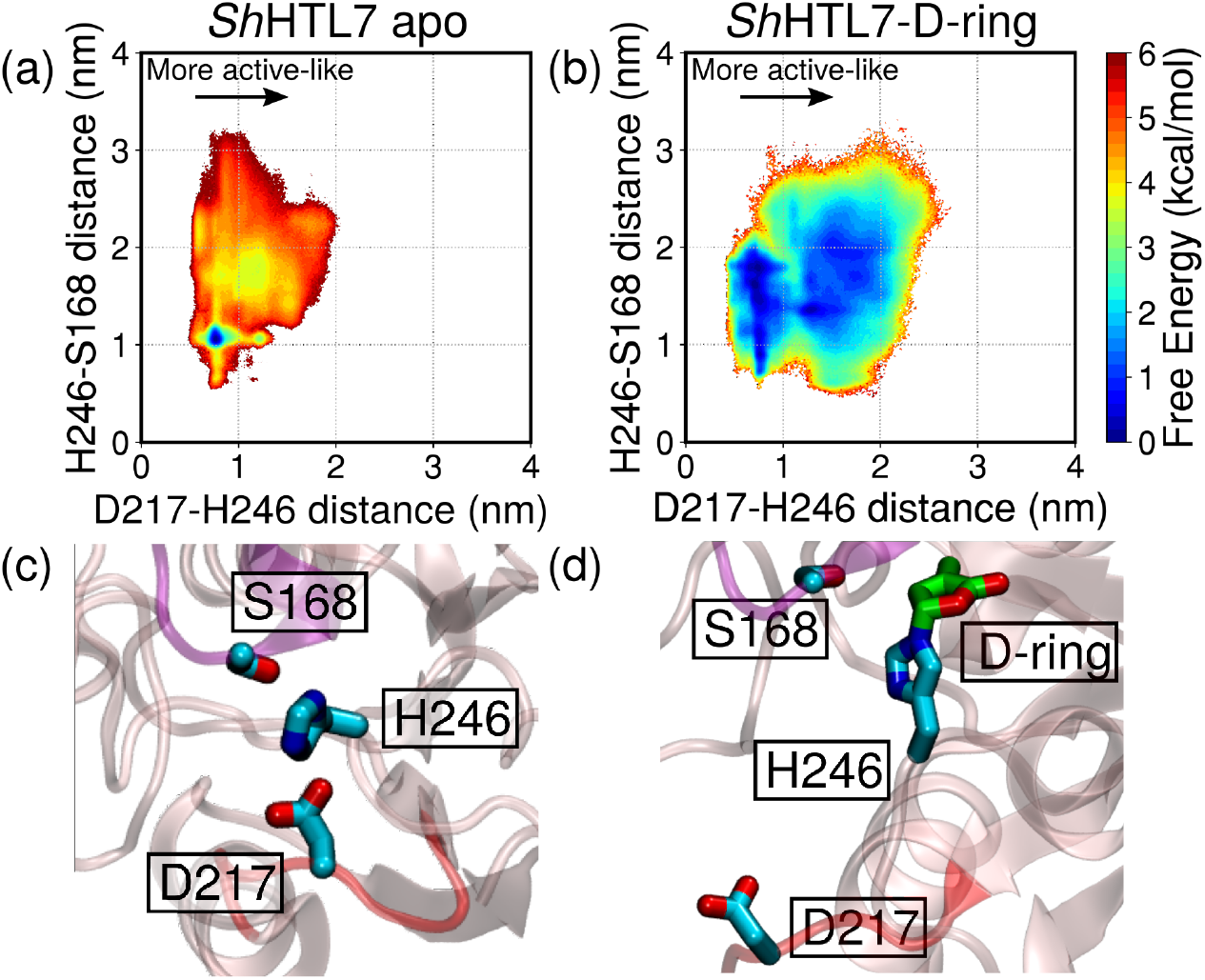
Free energy landscapes showing (a) the inhibition of D-loop detachment by the H246-S168 interaction in *apo Sh*HTL7 and (b) the disruption of the interaction in the presence of the covalent D-ring lowering the barrier for D-loop detachment. Images of the (c) S168-H246-D217 interaction and (d) disruption of the S168-H246 interaction by the covalent D-ring. More active-like states are found in regions with high D217-H246 distance, indicating detachment of the D-loop.

In addition to inhibiting the detachment of the D-loop, the S168-H246 contact in *apo Sh*HTL7 also inhibits the closure of the pocket via the T1-T3 helices. Disruption of this interaction increases the probability for these two residues to be far apart, which lowers the barrier for formation of active-like states in which the T1 and T3 helices are close together, as seen in Fig. 8a. This in turn promotes formation of T1-T3 closed conformations *Sh*HTL7 with the covalent D-ring over *apo Sh*HTL7. Another *apo Sh*HTL7-specific interaction that was found to inhibit T1-T3 closure, thus prevent full activation, is the H246-Q172 interaction (Fig. 8). In *apo Sh*HTL7, this interaction stabilized the receptor in partially active states in which T1 and T3 distances were insufficiently close together for full activation. The covalent D-ring disrupts this interaction, thus allowing the T1 and T3 helices to become close together and the receptor to undergo full activation (Fig. 8).

**Fig. 8.**
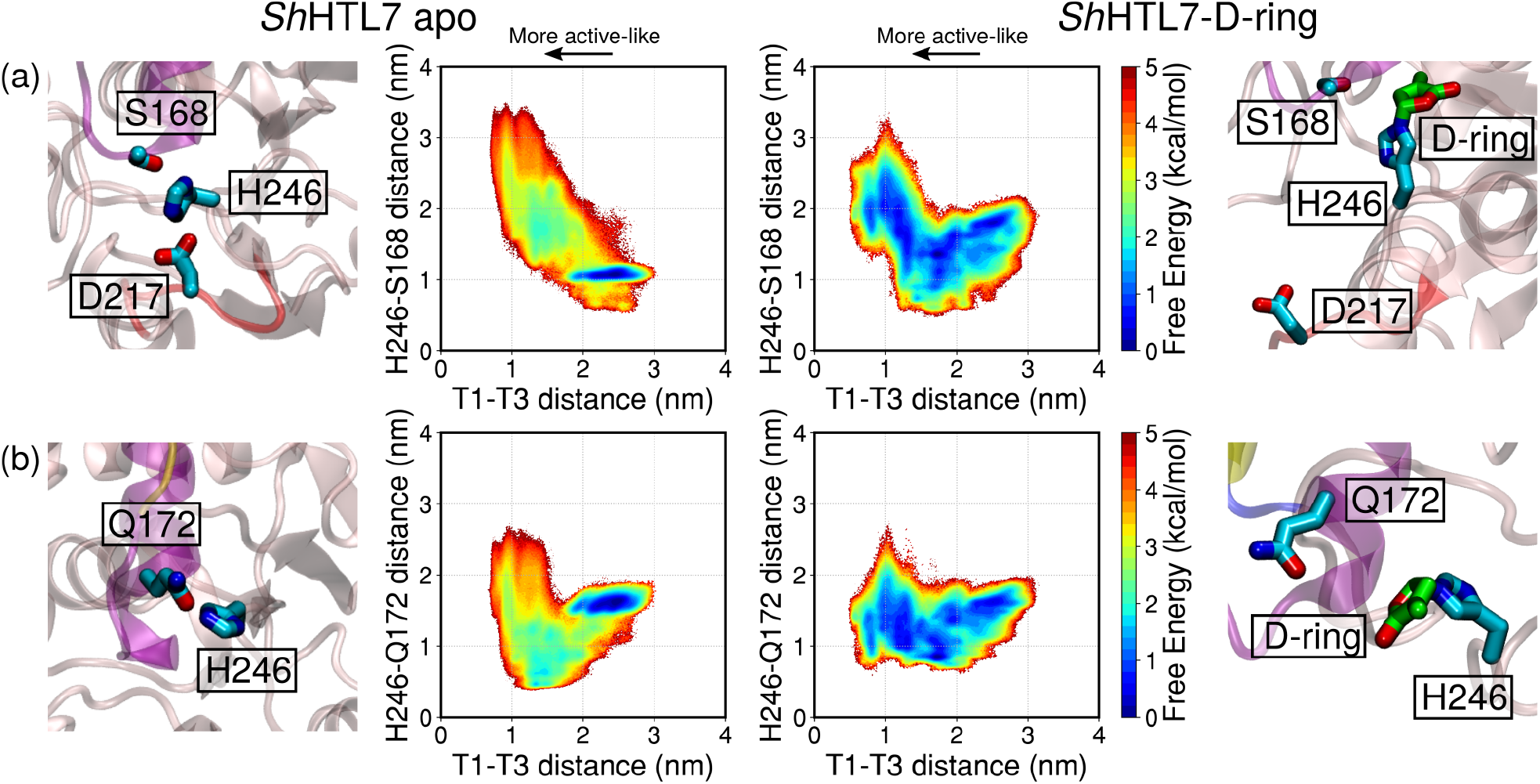
Free energy landscapes and accompanying structural images showing (a) the S168-H246 interaction that inhibits the activation of the T1-T3 closure molecular switch in *apo Sh*HTL7 and its disruption by the covalent D-ring and (b) the Q172-H246 interaction inhibiting T1-T3 closure in *apo Sh*HTL7 and its disruption by the covalent D-ring. More active-like states are found in regions with low T1-T3 distance, indicating closure of the binding pocket.

## Discussion

Using long-timescale molecular dynamics simulations with Markov state models, we have captured the activation pathways of *At* D14 and *Sh*HTL7 in the presence of a covalent butenolide ring (D-ring) attached to their catalytic histidine H247/246. In contrast to our previous results on the *apo* receptors,^15^ these simulations show that in the presence of the covalent D-ring, *Sh*HTL7 is more likely to transition to its active state than *At* D14. Since the butenolide modification is the result of strigolactone hydrolysis, this also means that *Sh*HTL7 has a greater response to the presence of strigolactone than *At* D14. This is consistent with observations that *Sh*HTL7 shows a high signaling response to the substrate despite its *in vitro* affinity for strigolactone being similar to only modestly higher than other strigolactone receptors.^10,11^

The necessity of strigolactone hydrolysis to induce signaling has been disputed by the existence of the catalytically inactive D218A mutant that is able to induce a downstream signaling response, ^16^ however, there has been evidence that hydrolysis acts as a promoter of signaling response either by formation of a covalently linked intermediate molecule (CLIM), which is an open butenolide (D-ring) bound to the catalytic serine and histidine^5^ or by formation of a covalent D-ring on the catalytic histidine. ^7^ In combination with our previous results on *apo* activation of strigolactone receptors, ^15^ we show that while *At* D14 and *Sh*HTL7 are both able to activate without substrate hydrolysis, the presence of the covalent D-ring resulting from strigolactone hydrolysis promotes activation. In both *At* D14 and *Sh*HTL7, key interactions between highly conserved residues and the covalent D-ring lower the free energy barrier of transitioning to the active state and stabilize the active state. Furthermore, in *Sh*HTL7, the covalent D-ring disrupts several *Sh*HTL7-specific H246 interactions that prevented the activation of key molecular switches. These H246 interactions were previously found to inhibit activation in *Sh*HTL7 but not *At* D14, which we proposed was a mechanism by which *Striga* evades suicidal germination. ^15^ Thus, it follows that disruption of these interactions leads to a much greater enhancement of activation in *Sh*HTL7 than in *At* D14 when the covalent D-ring is introduced. While the present work only considers the impact of the covalent D-ring on the histidine and not the CLIM that has been proposed to form, it is likely that a CLIM would have a similar effect on strigolactone receptor activation. Contact probabilities of *At* D14 and *Sh*HTL7 show that S97/95 has high contact probabilities with the covalent D-ring in addition to H247 (Fig. 2), so the CLIM is likely to form the same stabilizing interactions as the covalent D-ring. Additionally, the enhancement of activation in *Sh*HTL7 is largely modulated by disruption of contacts formed by H246 (Fig. 7-8), and since the CLIM would also be covalently bound to H246, these disruptions would also occur in the presence of CLIM. Finally, while there is disagreement whether CLIM or a covalent closed Dring is the driver of activation,^5,7,8^ a recent QM/MM study has suggested that both species are able to form during substrate hydrolysis, and that there are paths to interconversion between a CLIM and a closed D-ring on H247/246. ^29^

These results provide a mechanistic understanding of the process by which D-ringmodified *At* D14 and *Sh*HTL7 undergo conformational transition to their active state which is able to form a signaling complex with MAX2 protein to induce downstream effects. Notably, we also show that this activation process is a strong contributing factor to the high sensitivity of *Sh*HTL7 to striglactones, with D-ring-modified *Sh*HTL7 showing a ∼100-fold enhancement of activation over *apo Sh*HTL7, compared to only a ∼2-fold enhancement observed in *At* D14. Previously, we found that more efficient substrate binding and positioning for hydrolysis contribute about a ∼2-3 kcal/mol difference in the signaling pathways, which corresponds to roughly one or two orders of magnitude difference in signaling activity. ^27^The combined effects of substrate binding and receptor activation are thus able to account for roughly a ∼1000-10,000-fold difference in signaling activity between host and parasite strigolactone receptors. Since *Sh*HTL7 has been estimated to be ∼10,000-1,000,000 times more sensitive than other strigolactone receptors. ^10^ While the binding and activation steps are able to account for a significant portion of this difference, it is likely that other early steps in strigolactone signaling, in particular association with signaling partner MAX2, play a role in producing this differnce in sensitivity as well.

The insight that substrate hydrolysis-induced covalent modification enhances activation to a greater extent in *Sh*HTL7 than *At* D14 has important implications for the development of small-molecule modulators for witchweed control. Several small-molecule strigolactone receptor modulators have been proposed, including inhibitors^30–33^ and activators which induce suicidal germination in parasites.^34–36^ The insight that covalent D-ring modification potentently enhances activation in *Sh*HTL7 and to a much greater extent than in *At* D14 indicates that suicidal germination using compounds containing a butenolide moeity may be a potent and selective control strategy for witchweed. This is consistent with results reporting a femtomolar-affinity suicidal germination stimulant targeting *Sh*HTL7. ^35^

## Methods

### System Preparation

Active and inactive structures for *At* D14 were obtained from PDB codes 4IH4 and 5HZG, respectively. The inactive state structure of *Sh*HTL7 was obtained from PDB code 5Z7Y.^37^ The active state was constructed using a homology model using Modeller with 5HZG as a template.^5,13,14^ The structure of the covalent modfication was obtained from the proposed reaction mechanism.^5,7,8^ The protein was described using the AMBER ff14SB force field.^38^ The covalent modification to H247/246 was parameterized as a modified histidine residue. The histidine portion of the residue was described using ff14SB parameters, and the covalent ligand was described using GAFF.^39,40^ Each system was solvated in a ∼70×70×70 °A TIP3P water box with a 0.15 M NaCl concentration.

### Simulation Protocol

Each system was minimized for 10000 steps using a conjugate gradient descent algorithm followed by heating from 0 to 300K in the NVT ensemble and 10 ns equilibration at 300 K and 1.0 bar in the NPT ensemble. Temperature and pressure were maintained using the Langevin thermostat^41^ and Monte-Carlo barostat,^42^ respectively. Long-range electrostatics were computed using the particle mesh ewald algorithm. ^43^ An integration timestep of 2fs was used for all simulations. Bonds to hydrogen were constrained using the SHAKE algorithm. ^44^ All production simulations were performed using OpenMM 7.4.2. ^45^

### Adaptive Sampling

Rather than perform a single long molecular dynamics simulation to capture the activation process, an adaptive sampling scheme was used to sample the free energy landscape of activation. After each round of simulations, data was clustered on distances (molecular switches) associated with activation. Further simulations were seeded from the least sampled clusters in each round. Details of the adaptive sampling process including the number of trajectories and total simulation time for each round are included in Table S5 (*At* D14) and Table S6 (*Sh*HTL7). An aggregate of 805 *µ*s of simulations were performed on *At* D14, and an aggregate of 625 *µ*s of simulations were performed on *Sh*HTL7.

### Markov State Model Construction

Markov state models were constructed using PyEmma 2.5.4^46^ To select input features, 100 distances for *At* D14 and 60 distances for *Sh*HTL7 were chosen using the spectral oASIS method from the set of all lid domain distances.^47^ The full list of MSM input features is shown in Table S7 for *At* D14 and Table S8 for *Sh*HTL7. The distances were projected onto TICA components and discretized using the k-means algorithm. MSM hyperparameters (TICA components, number of clusters) were selected by maximizing VAMP scores (Fig. S5-6). MSM lag times were chosen by convergence of implied timescales with increasing lag time (Fig. S7). Final MSM hyperparameters are shown in Table S9. Validation of MSMs was performed using the Chapman-Kolmogorov test (Fig. S8). Markov state model probabilities were used to compute reweighted free energy landscapes using the relation in Eq. 1:

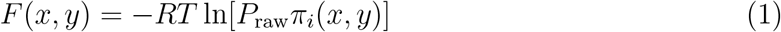

### Transition Path Theory

To resolve the top transition pathways from the active and inactive states of each receptor, we employed transition path theory (TPT).^28^ MSM clusters for each protein were lumped into five macrostates using PCCA clustering.^48^ Transition path theory was computed between the inactive and active macrostates using PyEmma 2.5.7^46^ and coarse grained onto the metastable set of macrostates. Top pathways that account for 99% of the total flux from the inactive to active states were identified.

### Molecular Switch Probability Calculations

Probabilities of each molecular switch being active were calculated by computing the activation probability each molecular switch within each MSM state, multiplying by the MSM equilibrium probability of each MSM state, and summing over MSM states as shown in Eq. 2.

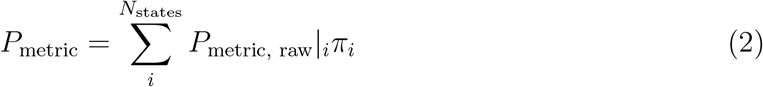

Overall probabilities of activation were computed using the product of individual molecular switch probabilities as shown in Eq. 3.

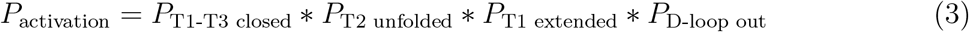

## Supporting information

Supplementary Methods, Images, Tables and Results

## Data and code availability

All in-house code used to analyze data and generate figures can be found at https://github.com/ShuklaGroup/Strigolactone-Covalent-Mod.

## Acknowlegments

This research was part of the Blue Waters sustained-petascale computing project, which was supported by the National Science Foundation (Award Nos. OCI-0725070 and ACI-1238993), the State of Illinois, and, as of December 2019, the National Geospatial-Intelligence Agency. Blue Waters was a joint effort of the University of Illinois at UrbanaChampaign and its National Center for Supercomputing Applications. The authors also acknowledge computational resources from the donors at the Folding@Home distributed computing platform and the Delta supercomputer at the National Center for Supercomputing Applications. J.C. is a member of the NIH Chemistry-Biology Interface Training Program (T32-GM136629). D.S. acknowledges support from the CAS Fellowship, Center for Advanced Studies at University of Illinois at UrbanaChampaign, and a Sloan Research Fellowship from the Alfred P. Sloan Foundation.

