## Supplementary Methods, Images, Tables and Results for "Effect of Histidine Covalent Modification on Strigolactone Receptor Activation and Selectivity"

### 1 Free energy landscapes projected onto TICA coordinates

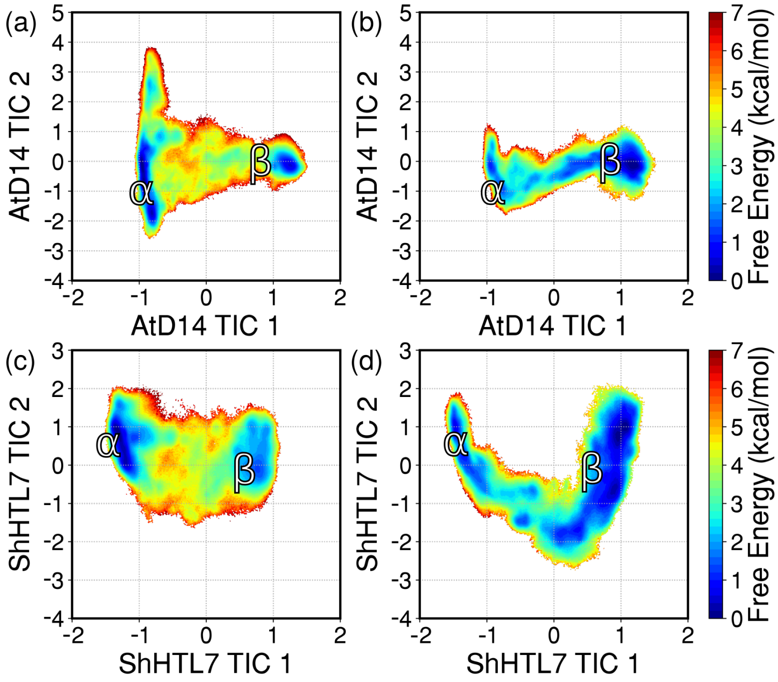


**Fig. S1.** Free energy landscapes of (a) *At*D14 simulation data projected onto *At*D14 TICA coordinates, (b) *Sh*HTL7 data projected onto *At*D14 TICA coordinates, (c) *At*D14 data projected onto *Sh*HTL7 TICA coordinates, and (d) *Sh*HTL7 data projected onto *Sh*HTL7 TICA coordinates. The inactive states are labeled as $\alpha$, and the active states are labeled as $\beta$.

### 2 Molecular switch and overall activation probabilities

| | *At*D14 *apo* | *At*D14-D-ring | *Sh*HTL7 *apo* | *Sh*HTL7-D-ring |
|---|---|---|---|---|
| T1-T3 closed | 0.356 | 0.274 | 0.037 | 0.354 |
| T1 extended | 0.316 | 0.399 | 0.192 | 0.439 |
| T2 unfolded | 0.324 | 0.574 | 0.276 | 0.643 |
| D-loop out | 0.308 | 0.366 | 0.242 | 0.583 |
| Overall activation probability | 0.011 | 0.023 | 0.0005 | 0.058 |

**Table S1.** Molecular switch probabilities for *apo* and covalent D-ring modified *At*D14 and *Sh*HTL7. These probabilities show a ∼2-fold overall enhancement of activation by the covalent modification in *At*D14 and a large ∼100-fold enhancement in *Sh*HTL7.

### 3 Top residues in contact with covalent D-ring

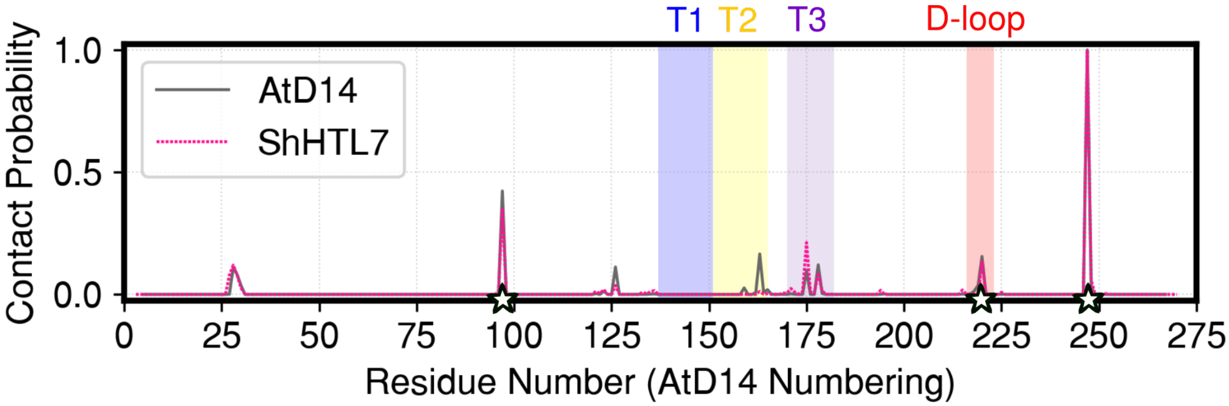


**Fig. S2.** Contact probabilities of each residue with the covalent D-ring. Residue numbering is sequence-aligned to *At*D14 numbering. Lid helices are labeled in blue, yellow, and purple, and the D-loop is labeled in red. Catalytic triad residues S97, D218, and H247 are labeled with stars.

| *At*D14 | | *Sh*HTL7 | |
|---|---|---|---|
| H247 | 1.000 | H246 | 1.000 |
| S97 | 0.423 | S95 | 0.354 |
| A163 | 0.166 | Y174 | 0.212 |
| S220 | 0.155 | M219 | 0.134 |
| T178 | 0.121 | Y26 | 0.120 |
| F28 | 0.112 | T177 | 0.088 |
| F126 | 0.112 | G25 | 0.078 |
| F175 | 0.099 | G27 | 0.078 |
| G29 | 0.080 | L247 | 0.057 |
| V219 | 0.037 | L124 | 0.036 |
| T30 | 0.036 | T28 | 0.025 |
| L248 | 0.029 | A170 | 0.024 |
| F159 | 0.027 | T121 | 0.020 |
| G165 | 0.019 | S214 | 0.018 |
| D218 | 0.014 | I193 | 0.017 |
| S123 | 0.013 | F134 | 0.015 |
| V164 | 0.008 | L178 | 0.015 |
| L162 | 0.008 | S119 | 0.013 |
| L179 | 0.004 | L161 | 0.012 |
| G121 | 0.003 | G133 | 0.012 |

**Table S2.** Residues with highest contact probability with the covalent D-ring in *At*D14 and *Sh*HTL7 and their respective contact probabilities. Residues for each protein are listed in descending order of covalent D-ring contact probability.

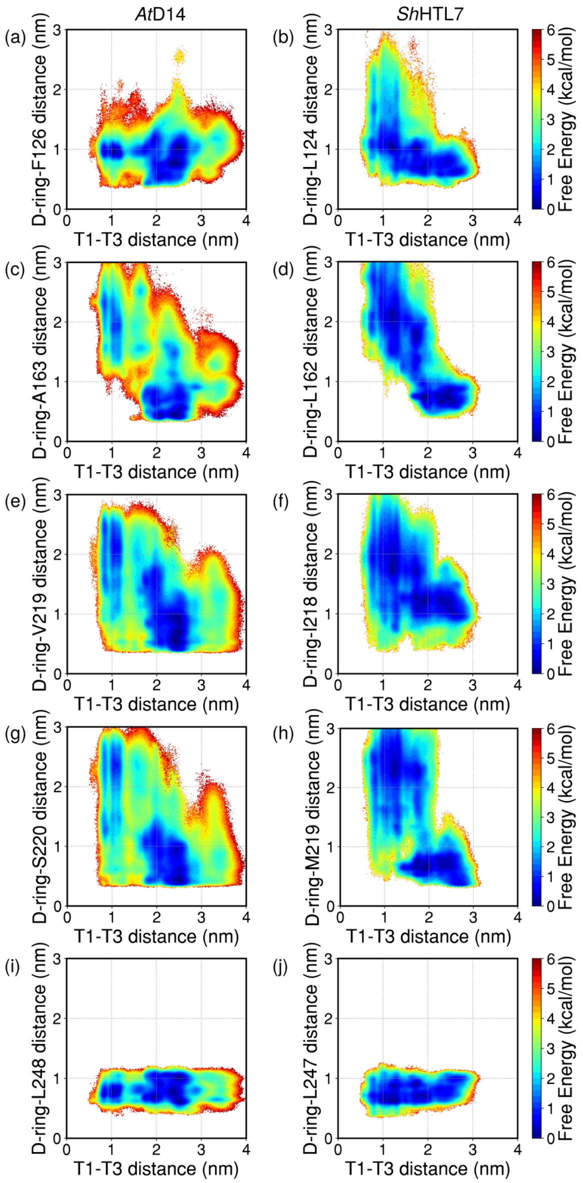


**Fig. S3.** Free energy landscapes projected onto T1-T3 distance and the distances between the covalent D-ring and F126/L124 (a,b), A163/L163 (c,d), V219/I218 (e,f), S220/M219 (g,h), and L248/L247 (i,j). These are covalent D-ring contacts that show high probability but little stabilization of the active state.

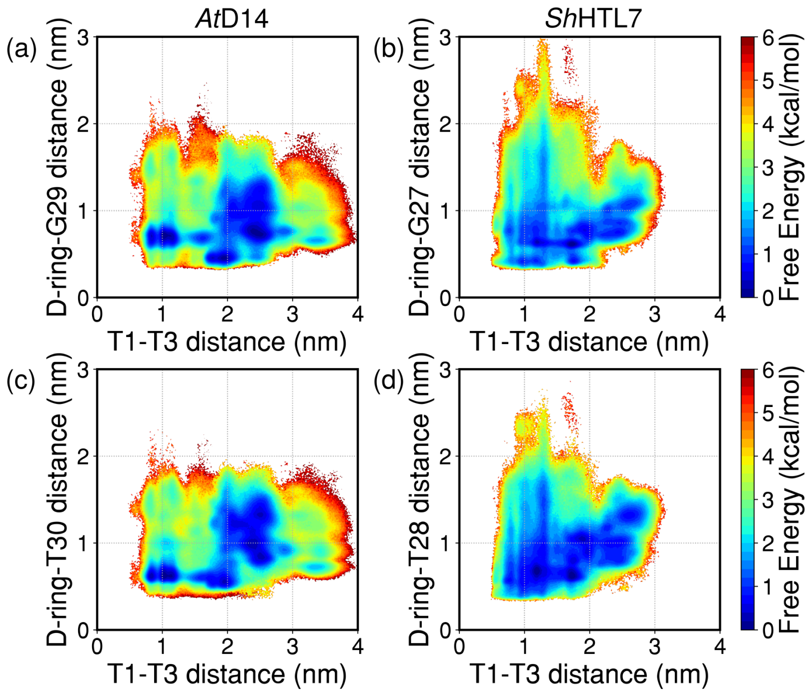


**Fig. S4.** Free energy landscapes projected onto T1-T3 distance and the distances between the covalent D-ring and G29/G27 (a,b) and the covalent D-ring and T30/T28 (c,d). Interactions of the covalent D-ring and these residues promote formation of the active state in a similar manner as interactions with F28/Y26 as shown in **Main Text**, Fig. 5.

### 4 Site conservation of key residues

| | *At*D14 | *Sh*HTL7 | *At*KAI2 |
|---|---|---|---|
| PDB ID | 4IH4 | 5Z7Y | 4HRX |
| Multiple Sequence Alignment | MAFFT | MAFFT | MAFFT |
| Homolog Source | UNIREF90 | UNIREF90 | UNIREF90 |
| Homolog Search Algorithm | HMMER | HMMER | HMMER |
| HMMER E-value | 0.0001 | 0.0001 | 0.0001 |
| HMMER Iterations | 1 | 1 | 1 |
| Maximal % Identity | 95 | 95 | 95 |
| Minimal % Identity | 50 | 50 | 50 |
| Number of Sequences | 150 | 150 | 150 |

**Table S3.** Parameters used for calculation of site conservation.

| Site | Conservation Score | Most Frequent Residues |
|---|---|---|
| *At*D14 Homologs | | |
| F28 | -0.574 | Y,F,L,V |
| G29 | -1.290 | G,C |
| T30 | -0.614 | M,I,S,C,A,G,T |
| F175 | -1.018 | L,V,I,C,F |
| T178 | -0.696 | S,H,T,Q,L,K,C,A,N,G,R |
| *Sh*HTL7 Homologs | | |
| Y26 | -0.419 | L,Y,F |
| G27 | -0.998 | G |
| T28 | -0.719 | T,S,M,C,V,I |
| Y174 | -0.478 | I,Y,F |
| T177 | -0.946 | T,M,A |
| *At*KAI2 Homologs | | |
| F26 | -0.797 | Y,F,L |
| G27 | -0.905 | G |
| T28 | -0.809 | T,S |
| F174 | -0.923 | F |
| T177 | -0.934 | T |

**Table S4.** Conservation of key sites in contact with the covalent D-ring in *At*D14, *Sh*HTL7, and *At*KAI2 and most frequent residues.

### 5 Summary of adaptive sampling rounds

Rather than perform single long simulations for each system, we applied an adaptive sampling scheme in which we ran multiple short simulations in parallel and seeded the next round of simulations from the least sampled conformations from the previous round of simulations. A table of adaptive sampling rounds is shown in Table S5 for *At*D14 and Table S6 for *Sh*HTL7. After the 17th adaptive round for each system, ~1500-2000 structures were selected from the aggregate data and run on the Folding@Home distributed computing platform. Subsequent adaptive rounds were seeded from the aggregate of rounds 1-17.

| *At*D14 | | | |
|---|---|---|---|
| Round | Parallel simulations | Approx. simulation time (ns) | Aggregate ($\mu$s) |
| 1 | 40 | 82 | 3.3 |
| 2 | 100 | 100 | 10.0 |
| 3 | 200 | 85 | 16.9 |
| 4 | 200 | 79 | 15.9 |
| 5 | 200 | 100 | 20.0 |
| 6 | 200 | 100 | 20.0 |
| 7 | 200 | 100 | 20.0 |
| 8 | 200 | 88 | 17.6 |
| 9 | 200 | 62 | 12.3 |
| 10 | 200 | 100 | 20.0 |
| 11 | 200 | 95 | 18.9 |
| 12 | 200 | 110 | 21.9 |
| 13 | 200 | 98 | 19.6 |
| 14 | 200 | 33 | 6.6 |
| 15 | 200 | 72 | 14.4 |
| 16 | 200 | 81 | 16.3 |
| 17* | 172 | 76 | 13.0 |
| 18 | 564 | 100 | 56.4 |
| 19 | 1000 | 100 | 100.0 |
| 20 | 471 | 100 | 47.1 |
| 21 | 161 | 100 | 16.1 |
| 22 | 211 | 100 | 21.1 |
| 23 | 200 | 100 | 20.0 |
| 24 | 300 | 100 | 30.0 |
| 25 | 497 | 100 | 49.7 |
| FAH | 2005 | 99 | 198.2 |
| | | Total ($\mu$s) | 805 |

**Table S5.** Summary of adaptive sampling rounds for *At*D14.

| *Sh*HTL7 | | | |
|---|---|---|---|
| Round | Parallel simulations | Approx. simulation time (ns) | Aggregate ($\mu$s) |
| 1 | 40 | 57 | 2.3 |
| 2 | 100 | 92 | 9.2 |
| 3 | 200 | 88 | 17.6 |
| 4 | 200 | 70 | 14.0 |
| 5 | 200 | 100 | 20.0 |
| 6 | 200 | 100 | 20.0 |
| 7 | 200 | 100 | 20.0 |
| 8 | 200 | 78 | 15.6 |
| 9 | 200 | 54 | 10.9 |
| 10 | 200 | 100 | 20.0 |
| 11 | 200 | 93 | 18.7 |
| 12 | 200 | 100 | 20.0 |
| 13 | 200 | 79 | 15.8 |
| 14 | 200 | 40 | 8.0 |
| 15 | 200 | 49 | 9.8 |
| 16 | 200 | 81 | 16.1 |
| 17* | 165 | 61 | 10.1 |
| 18 | 1500 | 100 | 150.0 |
| 19 | 500 | 100 | 50.0 |
| 20 | 280 | 100 | 28.0 |
| FAH | 1528 | 98 | 149.4 |
| | | Total ($\mu$s) | 625 |

**Table S6.** Summary of adaptive sampling rounds for *Sh*HTL7.

### 6 Distance features used for MSM construction

To construct Markov state models (MSMs), we computed 60-100 distance features which were selected using an automated approach (detailed in **Results of cross-validation tests used to select MSM hyperparameters**). The distances used for *At*D14 are listed in Table S7, and the distances used for *Sh*HTL7 are listed in Table S8.

| *At*D14 | | | |
|---|---|---|---|
| N128-P161 | F159-V219 | G139-V168 | G158-S224 |
| G165-S220 | A171-V219 | H157-V172 | E130-A170 |
| V164-K217 | E137-S220 | A163-N181 | H133-G246 |
| A166-P249 | D131-G158 | A170-G246 | N151-D167 |
| K143-A163 | A163-A223 | H157-S220 | G165-N181 |
| G158-E245 | G134-V219 | P222-E245 | D167-S224 |
| A150-G165 | D131-A166 | L162-P169 | E130-A223 |
| S220-L248 | D218-S224 | E153-A223 | H157-K217 |
| G158-A166 | A160-P222 | G165-R173 | F126-G134 |
| L162-G246 | P161-N181 | E137-A171 | E130-K217 |
| A171-P249 | P169-V221 | H133-S224 | D167-E245 |
| V164-A171 | E149-P169 | V156-V164 | A223-H247 |
| E153-D218 | A150-S224 | E130-A160 | G134-P222 |
| A163-D167 | E140-A166 | A147-R173 | D131-A171 |
| V156-P222 | D131-V164 | D167-N181 | M148-L162 |
| H157-A223 | P161-G165 | D131-V221 | E130-D167 |
| F159-R177 | H133-H157 | P161-S224 | G165-E245 |
| A166-A171 | A163-R173 | F175-V221 | G139-G158 |
| V172-D218 | F126-S220 | E140-V219 | E130-A150 |
| A154-V221 | A163-V219 | H157-P249 | V168-H247 |
| V156-V219 | L162-A166 | D131-P169 | P161-S220 |
| N181-A223 | V164-R177 | G135-D218 | A160-V168 |
| D218-G246 | M148-A160 | E138-S224 | G158-D218 |
| Y132-F159 | A150-P161 | L162-P222 | E153-A163 |
| R173-E245 | S224-P249 | G134-A170 | P169-K217 |

**Table S7.** All oASIS-selected features used for *At*D14 MSM construction. All inter-residue distances are C-$\alpha$ distances.

| *Sh*HTL7 | |
|---|---|
| D147-M181 | T157-M219 |
| E167-M219 | L160-G245 |
| E135-L161 | S154-E167 |
| L155-V222 | G156-R176 |
| K137-E167 | K151-M219 |
| E129-G156 | S145-A163 |
| S145-T157 | C164-L247 |
| P159-N180 | S154-D217 |
| E129-A169 | Y131-H246 |
| L153-H246 | L160-V220 |
| T157-M181 | A163-D217 |
| L155-P248 | A170-G245 |
| G132-D165 | P159-A170 |
| T157-D165 | Y174-M219 |
| E135-D217 | E148-L160 |
| S154-M181 | S152-A223 |
| G156-E244 | S154-V220 |
| L162-R176 | Y131-A158 |
| K137-L155 | L161-V222 |
| G132-E244 | S152-A158 |
| E129-M219 | K151-F179 |
| E148-P221 | F150-L162 |
| E167-E244 | L161-E167 |
| M181-M219 | E129-K151 |
| A163-A169 | A158-I218 |
| L124-A170 | E141-P159 |
| N180-N216 | E148-G156 |
| A163-F179 | V138-C164 |
| K151-E244 | E128-S154 |
| I218-G245 | L155-M219 |

**Table S8.** All oASIS-selected features used for *Sh*HTL7 MSM construction. All inter-residue distances are C-$\alpha$ distances.

### 7 Results of cross-validation tests used to select MSM hyperparameters

MSM hyperparameters were selected using GMRQ scores which were calculated using shuffle-split cross-validation. Features were automatically chosen using the spectral oASIS method with a feature set size determined by convergence of GMRQ with number of features. Following feature set selection, other hyperparameters were chosen by grid search with number of TICA components ranging from 2 to 8 and number of clusters ranging from 100 to 500. Selected parameters are shown in Table S9.

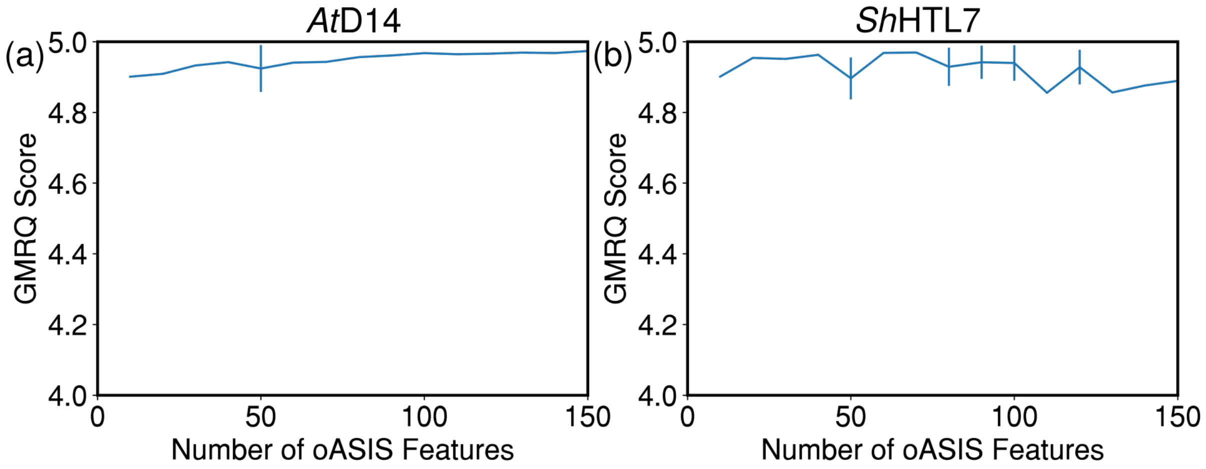


**Fig. S5.** Cross-validation scores of different sizes of oASIS feature sets. A feature set size of 100 was chosen for (a) *At*D14, and a feature set size of 60 was chosen for (b) *Sh*HTL7.

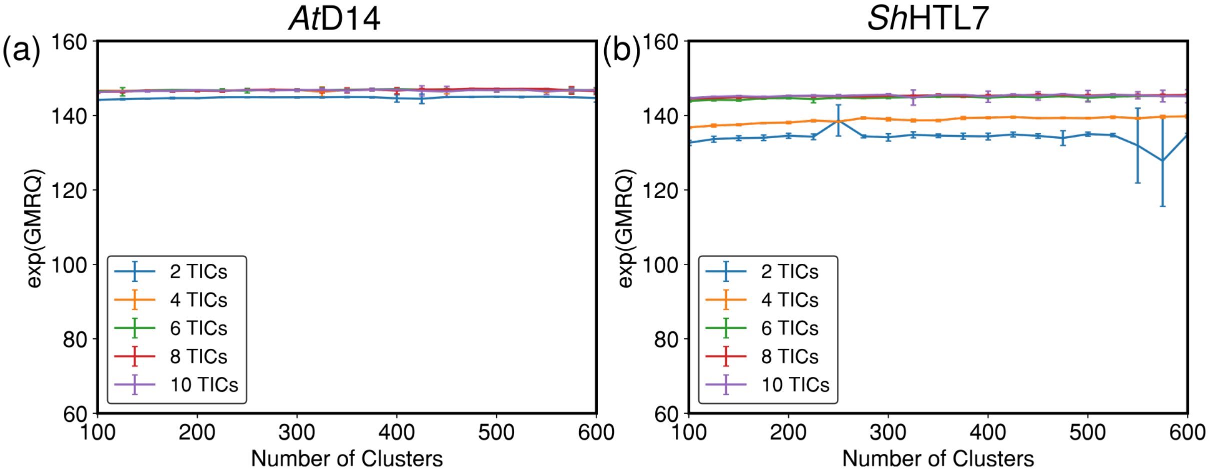


**Fig. S6.** Cross-validation scores with varying numbers of TICA components and clusters for (a) *At*D14 and (b) *Sh*HTL7.

### 8 Implied timescale plots used to select MSM lag times

MSM lag times were chosen using convergence of implied timescales. A lag time of 30 ns was chosen for both *At*D14 and *Sh*HTL7.

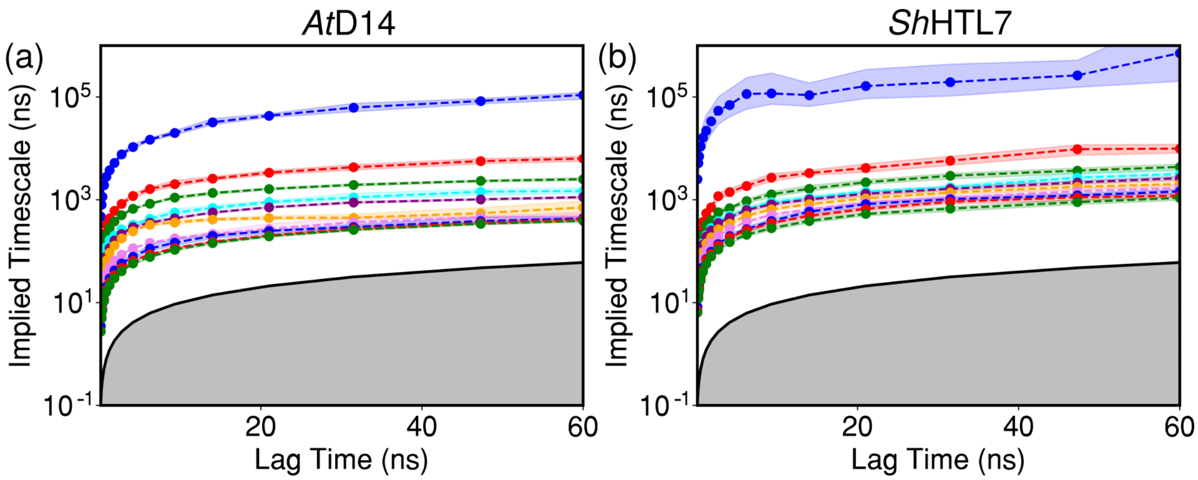


**Fig. S7.** Implied timescales of MSMs calculated at different lag times for (a) *At*D14 and (b) *Sh*HTL7.

| | $At$D14 | $Sh$HTL7 |
|---|---|---|
| Number of oASIS features | 100 | 60 |
| Lag time (ns) | 30 | 30 |
| Number of TICA components | 4 | 8 |
| Number of clusters | 375 | 350 |

**Table S9.** Final parameters used for MSM construction.

### 9 Chapman-Kolmogorov validation of MSMs

MSM validation was performed using the Chapman-Kolmogorov test. Briefly this tests for Markovianity of a system by comparing estimated transition matrices at lag times $n\tau$ at varying $n$ with predictions of the same transition matrices calculated at lag time $\tau$ multiplied by itself $n$ times.

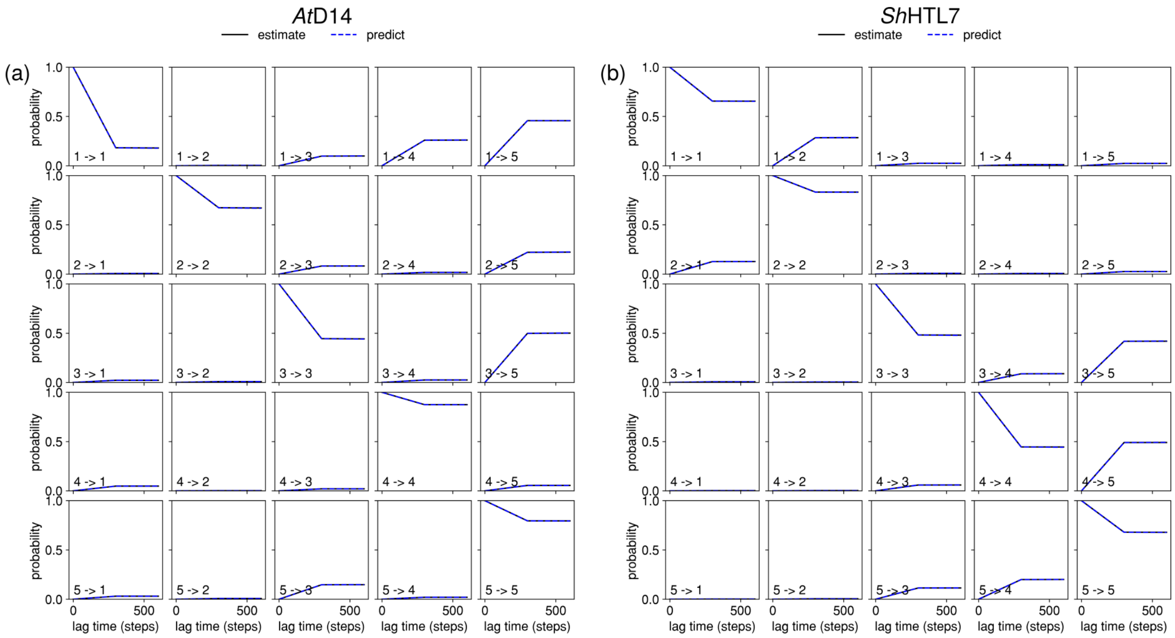


**Fig. S8.** Chapman-Kolmogorov validation for (a) $At$D14 and (b) $Sh$HTL7 MSMs.
